# Requirement for Fucosyltransferase 2 in Allergic Airway Hyperreactivity and Mucus Obstruction

**DOI:** 10.1101/2024.05.10.593580

**Authors:** Naoko Hara, Dorota S. Raclawska, Leslie E. Morgan, James C. NeeDell, Lucie Dao, Ayako Kato, Ana M. Jaramillo, Patrick S. Hume, Fernando Holguin, William J. Janssen, Eszter K. Vladar, Christopher M. Evans

## Abstract

Mucus hypersecretion is an important pathological problem in respiratory diseases. Mucus accumulates in the airways of people with asthma, and it contributes to airflow limitation by forming plugs that occlude airways. Current treatments have minimal effects on mucus or its chief components, the polymeric mucin glycoproteins MUC5AC and MUC5B. This treatment gap reflects a poor molecular understanding of mucins that could be used to determine how they contribute to airway obstruction. Due to the prominence of glycosylation as a defining characteristic of mucins, we investigated characteristics of mucin glycans in asthma and in a mouse model of allergic asthma. Mucin fucosylation was observed in asthma, and in healthy mice it was induced as part of a mucous metaplastic response to allergic inflammation. In allergically inflamed mouse airways, mucin fucosylation was dependent on the enzyme fucosyltransferase 2 (Fut2). *Fut2* gene deficient mice were protected from asthma-like airway hyperreactivity and mucus plugging. These findings provide mechanistic and translational links between observations in human asthma and a mouse model that may help improve therapeutic targeting of airway mucus.

## Introduction

Respiratory tissues are continuously exposed to inspired particles and potential pathogens whose accumulation can damage delicate gas exchange surfaces. Airway mucus is a critical first line of defense, but mucus can also be detrimental in lung diseases (1-5). In asthma, excessive or abnormal mucus is common across ranges of disease severity (2, 6), and airway mucus plugging is a causative feature in fatal asthma (5, 7). Despite its well-recognized associations with disease, effective therapies to prevent or reverse the effects of mucus overproduction in asthma are lacking. This treatment gap reflects an incomplete mechanistic understanding of mucus dysfunction at a molecular level (8).

The predominant macromolecules in airway mucus are the gel-forming mucin glycoproteins MUC5AC and MUC5B (8, 9). During synthesis, MUC5AC and MUC5B oligomerize through end-to-end disulfide bond formation, and they also become heavily glycosylated (10). The final products are massive glycopolymers with molecular weights in the mega-Dalton range.

Approximately 80% of mucin mass comprises carbohydrates that are added to long repetitive serine (Ser) and threonine (Thr) rich stretches of the protein while it traffics through the Golgi apparatus. Initial steps in mucin glycosylation involve addition of N-acetylgalactosamine (GalNAc) to Ser and Thr hydroxyls (Figure 1). N-acetylglucosamine (GlcNAc) and/or galactose (Gal) are then added to O-linked GalNAc to form Core glycan structures. Subsequent glycosylation steps include the addition of more GlcNAc and Gal glycans that form branches and extensions. Ultimately, glycosylation can be terminated by addition of sialic acid (Sia) or fucose (Fuc) to Gal.

**Figure 1.**
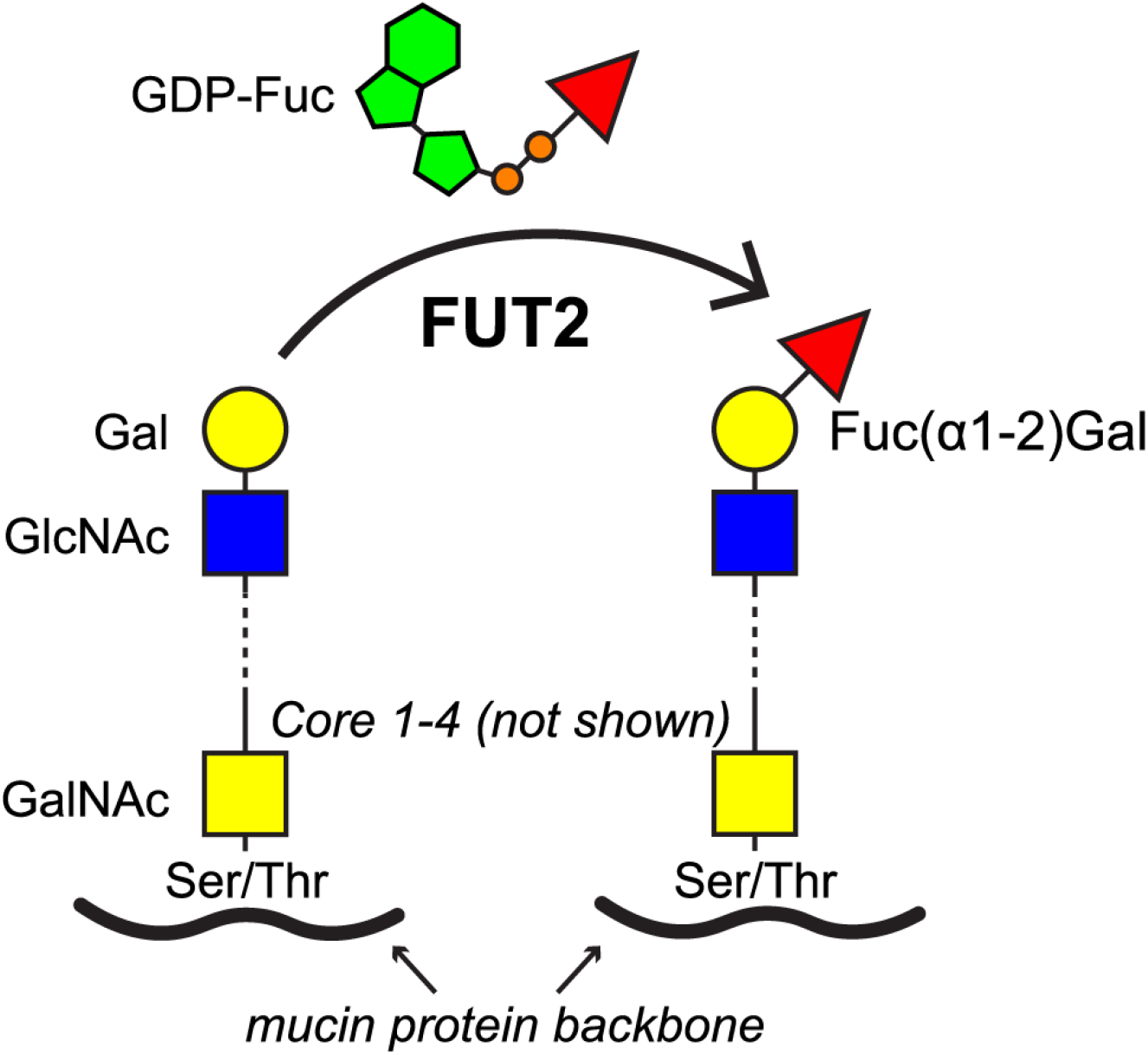
FUT2-mediated mucin fucosylation. Mucin fucosylation occurs via FUT2-mediated attachment of fucose (Fuc, red triangles) to galactose (Gal, yellow circles) glycans that extend from underlying (Ser) or threonine (Thr) residues possessing O-linked N-acetylgalactosamine (GalNAc, yellow squares) with or without N-acetylglucosamine (GlcNAc, blue squares) and Gal extensions. Fuc is transferred from a guanosine diphosphate (GDP)-linked carrier to Gal via bond formation between carbon 1 of Fuc and carbon 2 of Gal, resulting in a Fuc(α1-2)Gal structure.

Diversity in mucin glycosylation has evolved in part to mediate host interactions with microbes that recognize specific glycan structures. Although not necessarily mucin isoform specific, glycosylation varies across organs and even within anatomic sub-regions of tissues (11). Glycans on mucins also affect the biophysical properties of mucus gels due to effects on osmotic pressures exerted by the mass and charge of linked carbohydrates. Fuc is a mucin glycan whose polar surface area is approximately half of that presented by Sia. Accordingly, mucus that is rich in fucosylated mucin is less hydrophilic which may cause gels to be highly viscoelastic and prone to aggregation. While thick mucus may be helpful for host defense, it can lead to physiologic dysfunction. Indeed, fucose is suggested to be the most significant intrinsic determinant of abnormal viscoelasticity in chronic sinusitis mucus (12).

The addition of Fuc to Gal residues on mucins is mediated by the fucosyltransferase enzyme FUT2. FUT2 catalyzes formation of a glycosidic bond between the first carbon on Fuc and the second carbon on Gal thereby generating a Fuc(α1-2)Gal structure (hereafter referred to as α2-Fuc). This structure is also found on blood group secretor (or H) antigen, and H-positive secretor status increases asthma patients’ risks for severe exacerbations (13). We hypothesized mucus dysfunction may be explained in part by FUT2-mediated mucin α2-fucosylation. We tested this by investigating Fut2-mediated mucin fucosylation and its effects on airflow obstruction in a mouse model of allergic asthma.

## Methods

Detailed methods are provided in an online supplement.

### Human asthma and animal models

Fatal asthma lung samples used for histology were obtained from deidentified tissue donated to National Jewish Health. C57BL/6J congenic *Fut2*^−/−^ mice were generated previously (14) and were provided by Dr. Justin Sonnenburg at Stanford University. Males and females were used starting at 7-8 wks age. Mice were challenged using aerosolized *Aspergillus oryzae* extract (15) (AOE, 10% w/v, Cat. P6110, Sigma, St. Louis, MO) or saline vehicle administered weekly for 4 wks. Endpoints were collected 48 h after the last AOE challenge.

### Mucin detection

Triple labeling was performed to detect MUC5AC, MUC5B, and Fuc(α1-2)Gal in human and mouse airways. After antigen retrieval, tissues were blocked with Carbo-Free Blocking Solution (Vector Laboratories, Cat. SP-5040-125).

For human tissues, a mixture containing mouse-anti-human MUC5AC (clone 45M1, Novus, Cat. NBP2-15196, 1:1,000 dilution), rabbit-anti-human MUC5B (Sigma, Cat. HPA008246, 1:500 dilution), and biotin-conjugated *Ulex europaeus* agglutinin-1 (UEA1, Vector, B-1065-2, 8 μg/ml) was applied. UEA1 is selective for the glycan structure Fuc(α1-2)Gal (16).

Mouse mucins were detected using polyclonal rabbit-anti-mouse MUC5AC produced against the peptide N-CHALGDTSHAESSEQEFKSKESEEHGQQLAFR (Pacific Immunology, Ramona, CA, 1:2,500 dilution), mouse-anti-mouse MUC5B (clone MDA-3E1, Cat. MABT899, MilliporeSigma, Burlington, MA, 1:500 dilution), and UEA1 (as above).

Slides were cover-slipped with fluorescent mounting media containing DAPI. Negative controls included non-specific IgG, non-biotinylated UEA1, and knockout mice. Microscopy was performed using an Olympus BX63 microscope and cellSens software (Olympus USA, Center Valley, PA).

### Mucin glycoprotein blotting

For mouse lung lavage 5 × 0.5 ml saline was instilled into the right lungs. Aliquots of unmanipulated “ neat” lavage fluid were used for leukocyte enumeration or stored at -20° C. For blotting, 27 μl neat lavage samples were separated on 1% SDS agarose (100 V, 2 h), vacuum transferred to PVDF (17, 18), and blocked with Carbo-Free Blocking Solution (Vector). Mucins were probed with mixtures of UEA1 (1:1,000, 2 μg/ml) and either polyclonal rabbit-anti-mouse MUC5AC (1:2,000) or rabbit-anti-mouse MUC5B (1:5,000) (1, 19). Labels were detected with IRDye 680RD-conjugated streptavidin and IRDye 800CW-conjugated goat anti-rabbit IgG (1:15,000) using an Odyssey CLx and Image Studio software (LI-COR, Lincoln, NE).

### Airway hyperreactivity

As reported previously (15), lung function was measured using a flexiVent (Scireq, Montreal, Quebec, Canada). Mice were anesthetized with urethane (2.0 g/kg, i.p.), tracheostomized, ventilated (150 breaths/min, 10 ml/kg, 3 cm H_2_O positive end expiratory pressure) and paralyzed with succinylcholine (10 µg/g/min, i.p.). Total lung resistance (R_L_) and airway resistance (R_AW_) were determined at baseline and in response to nebulized methacholine (MCh; Sigma, St. Louis, MO; 0.1-10 mg/ml).

### Mucus plugging

During peak bronchoconstriction, mice were euthanized, and lungs were fixed as described previously (15, 18). Mucins were labeled using alcian blue-periodic acid Schiff’s (AB-PAS) stain (20). Total airway vs. parenchyma fractions were counted using a 250 × 250 µm grid, and airway mucin volume fractions were quantified using a 50 × 50 µm grid. Volume fractions and mucus volumes using were calculated.

### Statistical analyses

Statistical analyses were performed using GraphPad Prism 10.2.2 (397). Two-sample comparisons were compared using unpaired two-way *t*-tests or Mann-Whitney U-tests where appropriate. For multiple comparisons, ANOVA with a Dunnett post-hoc correction or a Kruskal-Wallis test with Dunn’s post-hoc correction was used.

## Results

### Mucins are α2-fucosylated in human asthmatic airways and in mouse models of allergic asthma

Mucus plugging is a prominent pathologic feature in fatal asthma (5). In a histologic sample from a fatal asthma subject, MUC5AC and MUC5B were prominent in goblet cells and in secreted mucus, which also demonstrated strong labeling for fucosylation with the lectin UEA1 (Figure 2A,B).

**Figure 2.**
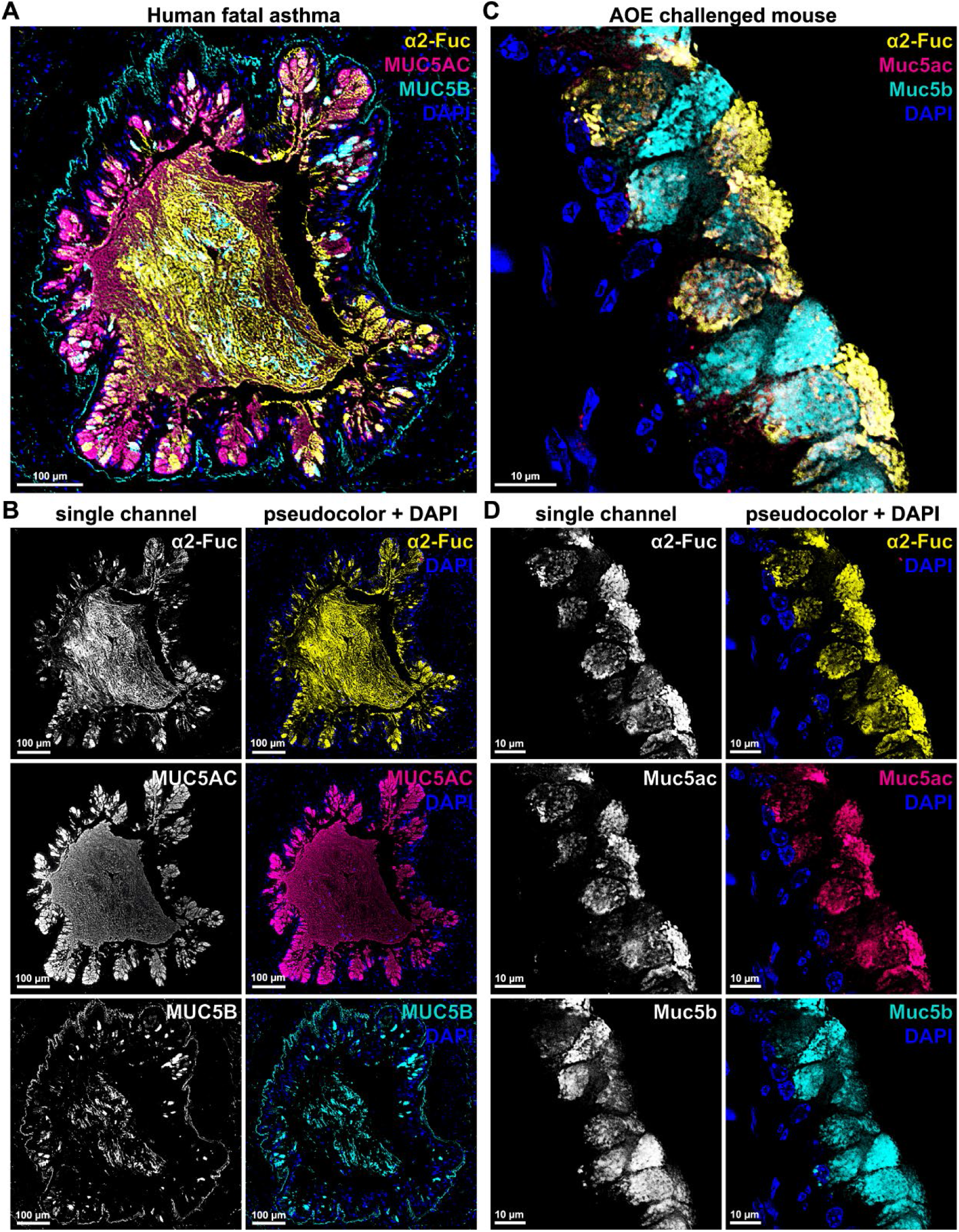
Fucosylation of airway polymeric mucins in human asthma and in a mouse model. **A,B**. In a human airway, α2-Fuc was detected using the lectin *Ulex europaeus* agglutinin-1 (UEA1, yellow, 8 μg/ml), which labeled airway mucous cells and a mucus plug in a person who died from fatal asthma. MUC5AC (magenta, 1:1,000) and MUC5B (cyan, 1:500) were both detected and fucosylated. **C,D**. In allergic mouse airways, Muc5b (cyan, 1:500), Muc5ac (magenta, 1:2,500) and α2-Fuc (UEA1, yellow, 8 μg/ml) labeling co-localize within airway epithelia. Monochrome (left) and pseudocolored individual label (right) versions are shown for human (**C**) and mouse (**D**) labels. Nuclei are stained with DAPI (blue).

Based on these findings, we assessed whether mucin fucosylation was also induced in a mouse model of allergic asthma. C57BL/6 mice were exposed to AOE and evaluated for mucin expression and α2-fucosylation histologically. Consistent with findings in human asthma, combined immunofluorescence and lectin labeling showed Muc5ac, Muc5b, and α2-Fuc in AOE-challenged mouse airway epithelia (Figure 2B,C), suggesting a role for Fut2 in mucin fucosylation.

To investigate a requirement for Fut2, we evaluated effects of *Fut2* gene disruption on airway mucin fucosylation in mice. In wild type (*Fut2*^+/+^) mice at baseline, frequent Muc5b-expressing cells were observed with little or no Muc5ac or α2-Fuc in the naïve state (Figure 3A). In airway epithelia of naïve *Fut2*^−/−^ mice, α2-Fuc was not detected, and steady-state Muc5b labeling was unchanged (Figure 3B). After AOE challenge, allergically inflamed mice showed robust induction of Muc5ac and α2-Fuc detection in *Fut2*^+/+^ airway epithelia (Figure 3C). *Fut2*^−/−^ mouse airways had undetectable UEA1 labeling after AOE, despite retaining Muc5b and Muc5ac expression (Figure 3D). These findings and a prior report showing a detrimental role for FUT2 in humans with asthma (13), prompted us to investigate whether Fut2 absence may protect mice from asthma-like airflow obstruction.

**Figure 3.**
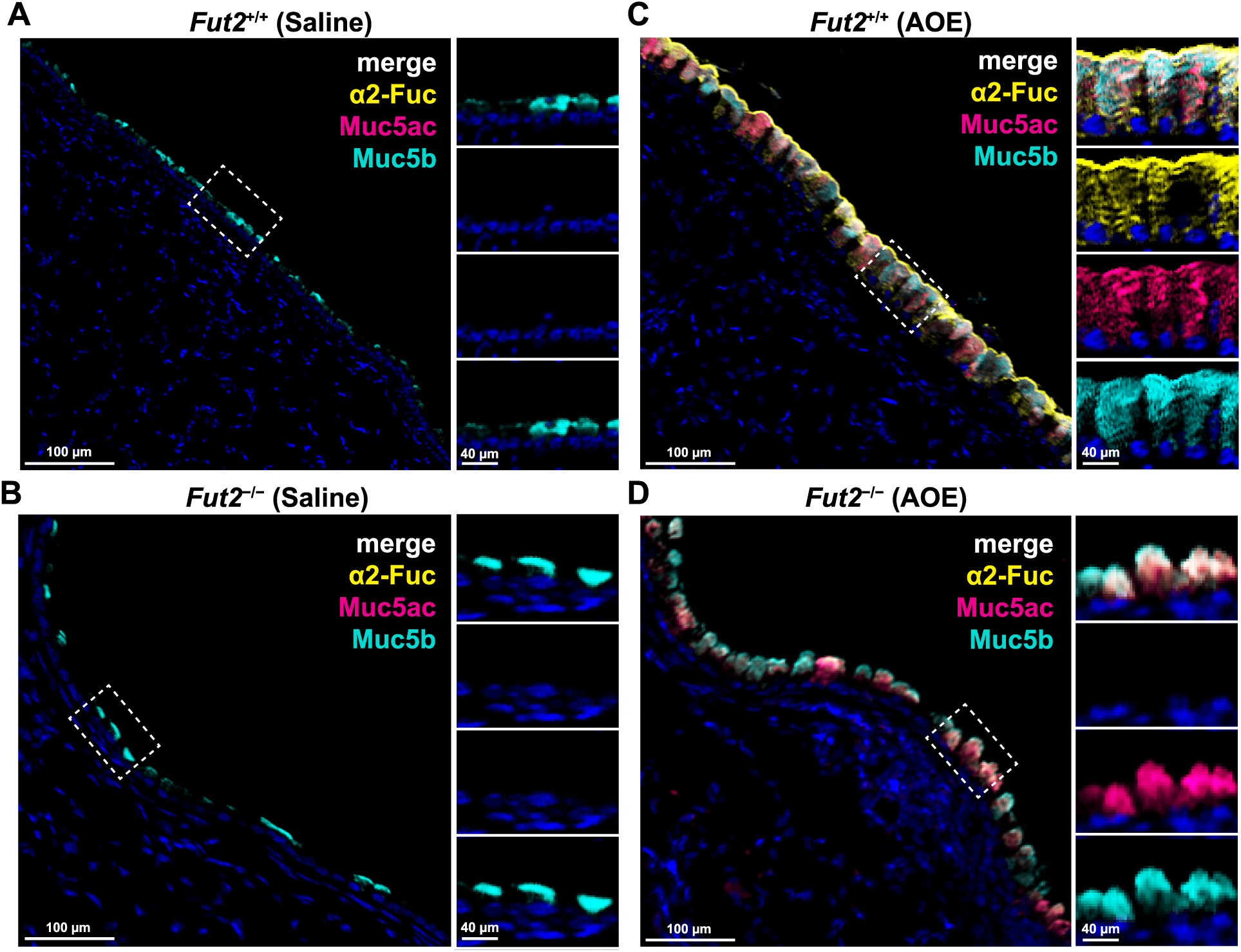
Allergic mouse airway epithelial α2-fucosylation requires *Fut2*. **A,B**. At baseline, α2-Fuc and Muc5ac are not detectable in airway epithelia in *Fut2*^+/+^ (**A**) of *Fut2*^−/−^ (**B**) mice, but Muc5b is observed. **C,D**. After AOE, Muc5b is sustained while Muc5ac and α2-Fuc detection are both induced in *Fut2*^+/+^ animals (**C**). In AOE-challenged *Fut2*^−/−^ mice, Muc5ac and Muc5b are sustained, but α2-Fuc detection is lost. Tissues were labeled with anti-Muc5b (cyan, 1:500), anti-Muc5ac (magenta, 1:2,500), biotinylated UEA1 (yellow, 8 μg/ml), and DAPI (blue).

### Fut2 is required for airway hyperreactivity in AOE challenged mice

We previously showed in AOE challenged mice that AHR can be abrogated by eliminating mucin from the airways (15, 18). To explore this further, we postulated that Fut2-dependent mucin fucosylation may play an essential role in causing muco-obstruction during AHR. In AOE challenged *Fut2*^+/+^ mice, airflow obstruction measured by dose dependent increases R_L_ and R_AW_ in response to MCh inhalation were potentiated compared to measurements in non-allergic controls. These characteristics of allergic asthma-like AHR were abolished in AOE challenged *Fut2*^−/−^ mice (Figure 4).

**Figure 4.**
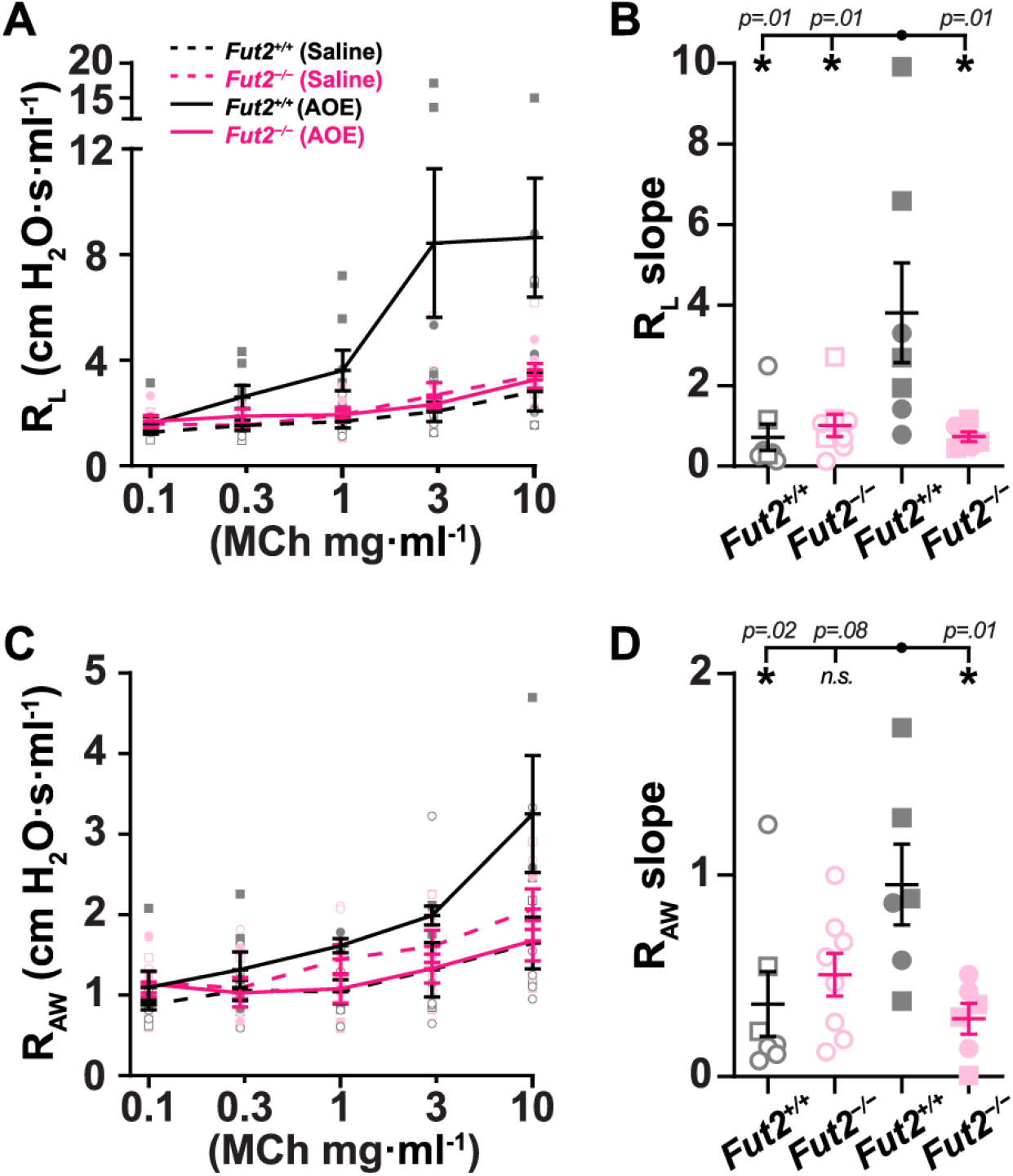
Fut2 is required for allergic airway hyperreactivity. Dose response curves to nebulized methacholine (MCh, 0.1-10 mg/ml) were generated in AOE and saline vehicle challenged *Fut2*^+/+^ (black) and *Fut2*^−/−^ (magenta) mice. Changes in total lung resistance (R_L_ in **A,B**) and conducting airway resistance (R_AW_ in **C,D**) in response to MCh were measured. Values are means ± sem with individual points per animal depicted by semi-transparent shapes. For each animal, dose response curves were fitted by log-linear best-fit regression analysis, and slopes of regression lines were compared by one-way ANOVA. ‘*’, p < 0.05 using Dunnett’s post-hoc test for multiple comparisons relative to AOE-challenged *Fut2*^+/+^ mice (p-values are shown in **B** and **D**). Comparisons were made between AOE challenged *Fut2*^+/+^ mice (filled gray, black solid lines, n = 7 biological replicates), AOE challenged *Fut2*^−/−^ mice (filled magenta, solid lines, n= 6 biological replicates), saline challenged *Fut2*^+/+^ mice (open gray, black dashed lines, n = 7 biological replicates for R_L_, n = 6 for R_AW_), and saline challenged *Fut2*^−/−^ mice (open magenta, dashed lines, n = 8 biological replicates). Squares identify males, and circles identify females.

Findings here are consistent with a previous report in a house dust mite (HDM)antigen challenge model of asthma (21). In that study, protection of *Fut2*^−/−^ mice from AHR was also associated with reduced allergic inflammation. To evaluate this, we compared changes in inflammation in AOE challenged *Fut2*^+/+^ and *Fut2*^−/−^ mice.

### Fut2 absence does not suppress allergic inflammation in AOE challenged mice

We enumerated leukocytes in lung lavage, and we observed that AOE strongly induced eosinophilic inflammation in both *Fut2*^+/+^ and *Fut2*^−/−^ mice (Figure 5, *see* Figure E1 in the data supplement). Importantly, numbers of leukocytes recovered in lavage fluid did not differ between mutant and wild type animals. To further validate these findings, we performed a multiplex cytokine analysis. Results showed few or no differences among pro-inflammatory and type 2 cytokines, or other markers (Tables E1,E2).

**Figure 5.**
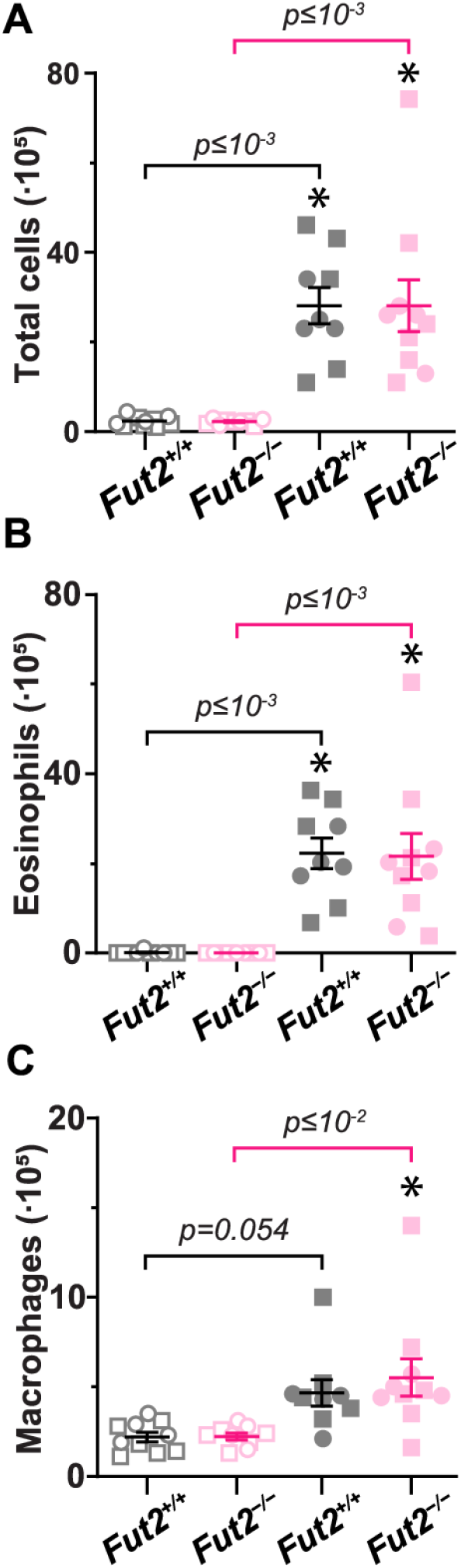
Fut2 deficiency but not reduce inflammation in AOE challenged mice. Lung lavage fluid was obtained from saline and AOE challenged *Fut2*^+/+^ (gray-black) and *Fut2*^−/−^ (magenta) mice. AOE challenge resulted in increases in total numbers of leukocytes (**A**), eosinophils (**B**) and macrophages (**C**) in both *Fut2*^+/+^ and *Fut2*^−/−^ mice compared to saline challenged controls. Kruskal-Wallis ANOVA was used to compare saline challenged *Fut2*^+/+^ mice (open gray shapes, n = 10 biological replicates), saline challenged *Fut2*^−/−^ mice (open magenta shapes, n = 9 biological replicates), AOE challenged *Fut2*^+/+^ mice (filled gray shapes, n = 9 biological replicates), and AOE challenged *Fut2*^−/−^ mice (filled magenta shapes, n = 10 biological replicates). ‘*’, p < 0.05 using Dunn’s post-hoc test for multiple comparisons (p-values are shown). Squares identify males, and circles identify females. No significant differences were observed using sex as a biological variable.

Protection from allergic AHR despite a lack of significant reduction inflammation was similar to prior observations in *Muc5ac* gene deficient animals (15). This suggested that protection was conferred by direct effects of Fut2 on mucins. Therefore, we performed immuno- and lectin blotting to evaluate mucins in lung lavage samples collected from *Fut2*^+/+^ and *Fut2*^−/−^ mice.

Irrespective of genotype, both Muc5ac and α2-Fuc detection were low in the Muc5b-rich samples collected from naïve mice (Figures 6 and E2). After AOE challenge, there was a strong induction of mucin α2-fucosylation in lavage samples from *Fut2*^+/+^ mice. Muc5b and Muc5ac were both increased and α2-fucosylated. Colocalization of UEA1 signals validated that mucins themselves carry α2-Fuc glycans after AOE-induced allergic inflammation (Figures 6 and E2).

**Figure 6.**
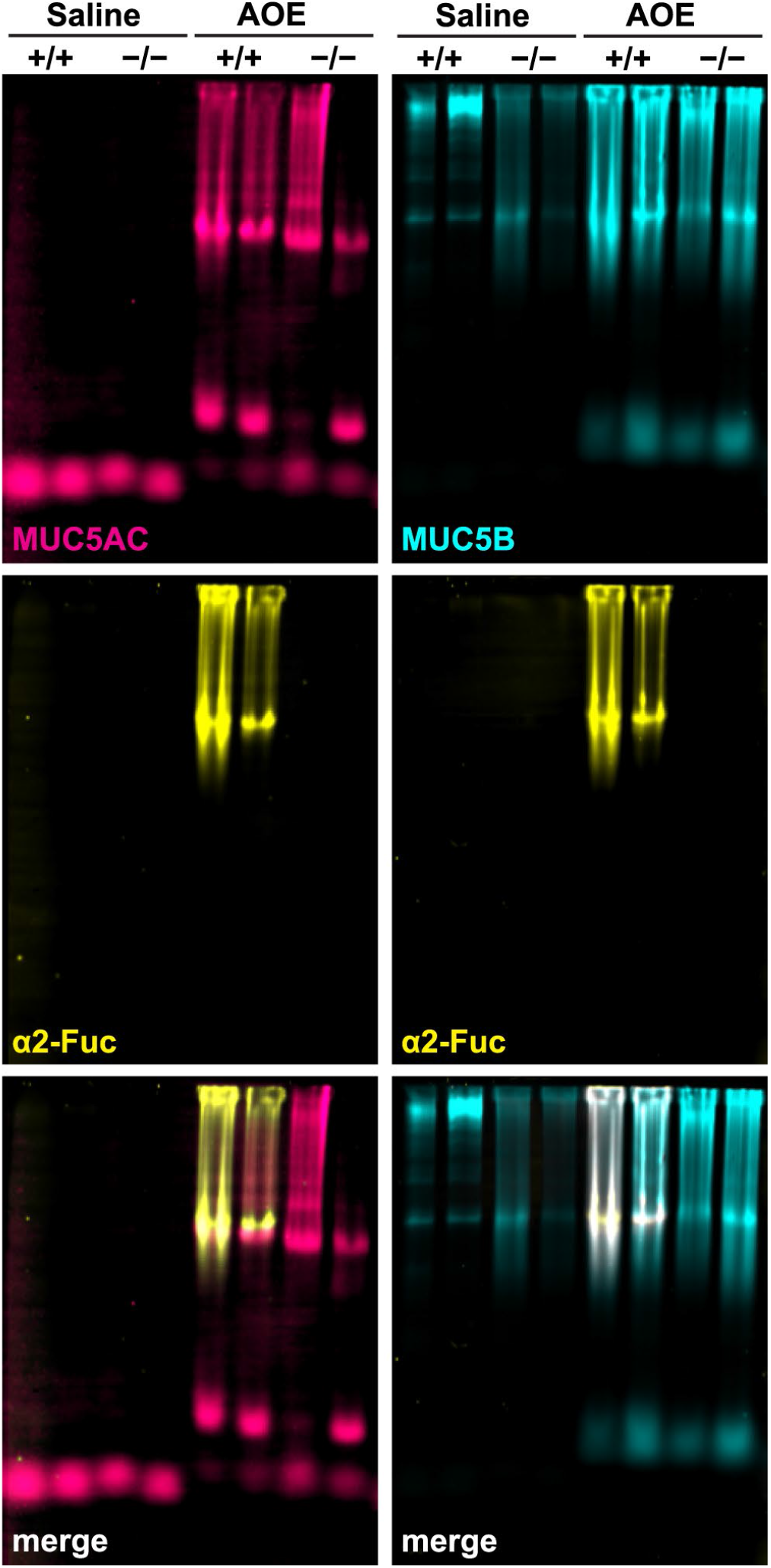
Fut2 is required for Muc5ac and Muc5b α2-fucosylation. Combined immuno- and lectin blotting was performed on neat lavage. Equal volumes of lavage fluid (27 μl) were loaded per well, separated on 1% SDS agarose gels, and transferred to PVDF membranes. Membranes were probed with biotinylated UEA1 (1:1,000, 2 μg/ml) and either rabbit-anti-Muc5ac (1:2,000) or rabbit-anti-Muc5b (1:5,000). For secondary detection, Alexa 680-conjugated streptavidin and Alexa 800-conjugated labeled goat-anti-rabbit probes (Licor, 1:15,000) were applied. Monochrome images were acquired and pseudocolored magenta (Muc5ac), cyan (Muc5b), or yellow (α2-Fuc). Image overlays demonstrate UEA1 and mucin signal colocalization (white in merged panels).

By contrast, in samples collected from AOE challenged *Fut2*^−/−^ mice, α2-Fuc detection was absent even though Muc5ac and Muc5b were both still present. Since fucosylation increases mucus viscoelasticity (12), and *Fut2*^−/−^ mice were protected from AHR (see Figure 4), we hypothesized that Fut2 potentiates allergic AHR by promoting mucus obstruction.

### Fut2 promotes airway mucus obstruction

We assessed secreted mucus volume, airway occlusion, and heterogeneous plugging in AOE challenged mice during MCh-induced bronchoconstriction. In *Fut2*^+/+^ mouse airways, secreted mucus aggregated on airway surfaces (Figure 7A), but accumulated mucus was less apparent in airways from *Fut2*^−/−^ mice (Figure 7B). These observations were validated by stereological quantification. Bronchial mucus volumes and fractional mucin occlusion were significantly decreased in *Fut2*^−/−^ mice compared to *Fut2*^+/+^ animals (Figure 7C,D and E3).

**Figure 7.**
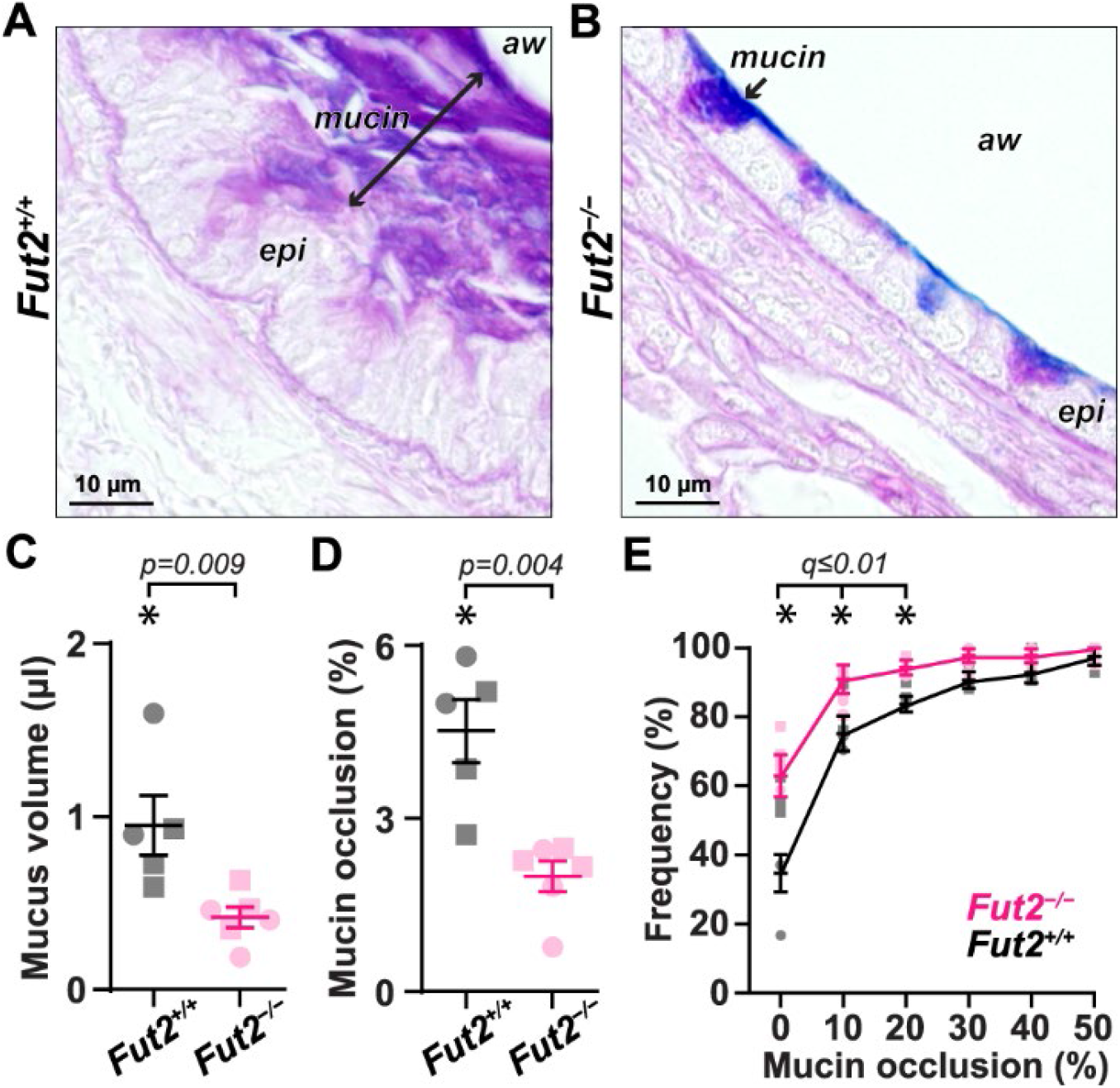
Fut2 deficiency reduces airway mucus plugging. After lung mechanics studies, lungs were fixed with methacarn to preserve mucus in airspaces. **A,B**. AB-PAS stained tissues of *Fut2*^+/+^ mice show mucin aggregated on airway surfaces (**A**) that was less prominent in *Fut2*^−/−^ animals (**B**); ‘epi’ is epithelium, and ‘aw’ is airway lumen. **C-E**. Calculated mucus volumes (**C**), fractional mucin occlusion (**D**), and heterogeneous plugging (**E**) were significantly decreased in bronchial airways of AOE challenged *Fut2*^−/−^ mice. Data in **C** and **D** are means ± sem for *Fut2*^+/+^ (gray shapes, black lines, n = 5 mice) and *Fut2*^−/−^ (magenta shapes and lines, n = 6 mice). P-values are shown with ‘*’ depicting p<0.05 by two-tailed Mann-Whitney tests. Data in **E** are means ± sem displayed on cumulative frequency distributions. Fractional occlusion means per animal were evaluated between *Fut2*^+/+^ and *Fut2*^−/−^ mice using multiple unpaired t-tests with two-stage step-up methods for multiple comparisons at a false discovery rate cut-off (q-value) set at 0.05. ‘*’ demonstrates significance, and q-values are shown in **E**. In **C-E**, semi-transparent shapes identify results from individual mice, squares identify males, and circles identify females. No significant differences were observed using sex as a biological variable.

Overall mean levels of mucus obstruction do not fully reflect the heterogenous nature of mucus obstruction that drives AHR (15, 18). We thus evaluated frequencies of airway occlusion at different magnitudes across airway levels. Consistent with prior findings in Muc5ac deficient and mucolytic treated mice (15, 18), AOE challenged *Fut2*^−/−^ mice demonstrated significantly greater proportions of airways with 20% or less occlusion (Figure 7E). Thus, Fut2 absence in mice resulted in protection from a heterogeneous muco-obstructive phenotype similar to what is seen in asthma (see Figure 2) (5).

## Discussion

The studies reported here demonstrate that mucin fucosylation is associated with allergic mucous metaplasia (see Figure 2), and it plays a causative role in asthma-like airflow obstruction in mice. A role for FUT2 is supported by findings in *Fut2* gene disrupted mice that lacked fucosylated airway mucins and are protected from AHR and airway mucus plugging even as allergic inflammation persisted (see Figures 3-7). These findings provide translational and causative links to prior reports showing associations between mucin fucosylation in human asthma and animal models (13, 22, 23)

*FUT2* and mucin gene loci are associated with chronic sputum production (24), and people with asthma who lack FUT2 are protected from severe exacerbations (13). Over extended periods, these could be explained by effects on host defense and inflammation that are complex and are also both organism and tissue specific. Indeed, *FUT2* gene status can affect how hosts respond to viral, bacterial, and fungal microorganisms in different organs.

People with *FUT2* loss-of-function mutations are protected from viral infections including HIV (25, 26), norovirus (27), and rotavirus (28) that each use Fuc(α1-2)Gal carbohydrate linkages for host cell attachment. FUT2 absence is also associated with bacterial infections in the middle ear (29) and the lungs (30), and with the dysbiosis and pathogen infection in the gastrointestinal and female reproductive tracts (31-33). In the intestine, FUT2 helps shape the gut microbiota (31), type 3 innate lymphoid cells regulate its expression (32), and potentially immune tone in distant organs such as the lung.

A connection between Fut2 and the development of allergic inflammation was previously explored in an HDM antigen exposure model (21). Similar to what is reported here, HDM-challenged *Fut2* deficient mice were protected from AHR, and their airways were devoid of UEA1-positive signals. That report also found partial reductions in inflammatory markers including eosinophils in lung lavage, type 2 and 17 lymphocytes and monocyte-derived dendritic cells in lung tissues, and IgG1 specific for HDM antigens in serum. However, the timepoint used for assessing inflammation and AHR was day 12 after the start of sensitization (21), suggesting a potential role for Fut2 in processes related to the development of allergy to HDM.

We did not observe a decrease in inflammation in mice challenged with AOE (see Figures 4 and E3). Our disparate findings could reflect differences between HDM and AOE as allergens. Furthermore, our choice of a more distant endpoint (23 days after the first AOE challenge) may have allowed allergic sensitization to develop more completely in both wild type and knockout strains. Lavage leukocytes increased in AOE challenged animals irrespective of *Fut2* genotype (Figures 5 and E1), and there were few differences in cytokine profiles (Tables E1,2). Accordingly, inflammosuppressive effects of FUT2 absence do not account for protection from AHR in findings reported here.

Fungal derived fucose-selective lectins may reveal relevant relationships between type 2 immune signals and the upregulation of mucin fucosylation in the lungs. *Aspergillus fumigatus* expresses a fucose-specific lectin called FleA that binds to mucins (34, 35). *Aspergillus oryzae* expresses a FleA lectin as well (36). FleA-mucin interactions are proposed as important means through which fucosylated mucins support innate host defense by trapping fungal conidia and allowing their elimination by mucociliary clearance (34, 35). This induction of a fucosylated mucous phenotype is consistent with aberrant overlaps observed between protective type 2 immunity and allergy.

Without a need for type 2 mediated immune defense, the acquisition of a heavily fucosylated mucin glycotype in allergically inflamed lungs results in mucus dysfunction. Fucose is a low charge glycan that increases mucus viscoelasticity (12). An excessively viscoelastic mucus gel can overwhelm mucociliary clearance or even cough-generated forces. In lower airways, the production of MUC5AC and a low-charge glycoform of MUC5B is associated with altered mucin composition in asthma (37, 38), and this is a prominent finding in fatal cases (39). In addition to intrinsic viscoelastic properties conferred by α2-Fuc, it is also possible that lectins from *A. oryzae* or other species could bind fucosylated mucins and cause aggregation as well. It is thus plausible that, in addition to the acute mucus plugging observed in response to MCh, persistent mucus aggregation caused by fungal lectins could have chronically detrimental effects in patients with allergic asthma. Each of these associations could be explained in part as resulting from excessive fucosylation (40) (see Figure 2A), but additional mechanistic work is needed.

Based on our findings, two important next steps are to determine mechanisms that regulate FUT2 expression and function. Expression studies in mice suggest tissue specific programs. In the gastrointestinal tract, IL-22 drives Fut2-mediated fucosylation phenotypes (32). However, IL-22 is dispensable for α2-fucosylation in the airways where *Fut2* gene expression and fucosylation appear to be mediated predominantly in an IL-13/STAT6-dependent manner. Notably, *Fut2* expression and α2-fucosylation in the airways are retained in *Rag2*^-/-^ mice, suggesting a direct effect of IL-13/STAT6 signaling in airway epithelia (21).

It is also likely that differences in inflammatory and transcriptional programs that regulate mucin fucosylation reflect genetic variation among (and within) species. Associations of *FUT2* loss-of-function mutations and mucin gene loci with chronic sputum production were found in people even without prior respiratory disease diagnoses (24). More work is needed to clarify gene regulation mechanisms that drive tissue specific *Fut2* expression.

Independent of upstream pathways that regulate *FUT2* expression, there is also a need to better understand FUT2 enzyme activity. Although little is known about FUT2, high-resolution structural characterizations of FUT8 and FUT9 have been made (41-45). They reveal crucial sites of GDP-fucose docking (see Figure 1) and catalysis of Fuc(α1-6)GlcNAc and Fuc(α1-3)GlcNAc bond formation. They also identify allosteric sites on FUT8 that mediate its activity, especially with respect to its function in the low pH environment of the Golgi apparatus (42, 43, 45).

In summary, mucus obstruction plays pathological roles in asthma and other pulmonary diseases. Presently, there are no effective drugs that specifically target mucins. The findings reported here demonstrate a detrimental role for FUT2-mediated fucosylation in mucus obstruction. FUT2 inhibition is presently limited to utilization of non-selective modified fucose derivatives (46), but FUT2 selective antagonists are being developed for treating cancer (47). A deeper understanding of *FUT2* gene regulation and FUT2 structure-function relationships may guide development of specific inhibitors.

## Acknowledgement of Funding Support

NIH R01HL080396, R01HL130938, P01HL162607, K99HL165072, R01HL14974, R35HL140039, K08HL155894, and VA Merit I01BX005343.

## References

1. Roy MG, Livraghi-Butrico A, Fletcher AA, McElwee MM, Evans SE, Boerner RM, et al. Muc5b is required for airway defence. Nature. 2014;505(7483):412–6. Epub 2013/12/10. doi: 10.1038/nature12807. PubMed PMID: 24317696; PubMed Central PMCID: 4001806.

2. Dunican EM, Elicker BM, Gierada DS, Nagle SK, Schiebler ML, Newell JD, et al. Mucus plugs in patients with asthma linked to eosinophilia and airflow obstruction. The Journal of clinical investigation. 2018;128(3):997–1009. Epub 2018/02/06. doi: 10.1172/JCI95693. PubMed PMID: 29400693; PubMed Central PMCID: 5824874.

3. O’Riordan TG, Zwang J, Smaldone GC. Mucociliary clearance in adult asthma. The American review of respiratory disease. 1992;146(3):598–603. Epub 1992/09/01. doi: 10.1164/ajrccm/146.3.598. PubMed PMID: 1519834.

4. Messina MS, O’Riordan TG, Smaldone GC. Changes in mucociliary clearance during acute exacerbations of asthma. The American review of respiratory disease. 1991;143(5 Pt 1):993–7. Epub 1991/05/01. doi: 10.1164/ajrccm/143.5_Pt_1.993. PubMed PMID: 2024856.

5. Kuyper LM, Pare PD, Hogg JC, Lambert RK, Ionescu D, Woods R, et al. Characterization of airway plugging in fatal asthma. AmJMed. 2003;115(1):6–11. PubMed PMID: 12867228.

6. Ordonez CL, Khashayar R, Wong HH, Ferrando R, Wu R, Hyde DM, et al. Mild and moderate asthma is associated with airway goblet cell hyperplasia and abnormalities in mucin gene expression. American journal of respiratory and critical care medicine. 2001;163(2):517–23. Epub 2001/02/17. doi: 10.1164/ajrccm.163.2.2004039. PubMed PMID: 11179133.

7. Dunnill MS. The pathology of asthma, with special reference to changes in the bronchial mucosa. JClinPathol. 1960;13:27–33. PubMed PMID: 13818688.

8. Fahy JV, Dickey BF. Airway mucus function and dysfunction. The New England journal of medicine. 2010;363(23):2233–47. Epub 2010/12/03. doi: 10.1056/NEJMra0910061. PubMed PMID: 21121836; PubMed Central PMCID: 4048736.

9. Boucher RC. Muco-Obstructive Lung Diseases. The New England journal of medicine. 2019;380(20):1941–53. Epub 2019/05/16. doi: 10.1056/NEJMra1813799. PubMed PMID: 31091375.

10. McShane A, Bath J, Jaramillo AM, Ridley C, Walsh AA, Evans CM, et al. Mucus. Current biology : CB. 2021;31(15):R938–R45. doi: 10.1016/j.cub.2021.06.093. PubMed PMID: 34375594; PubMed Central PMCID: PMC8759706.

11. Becker DJ, Lowe JB. Fucose: biosynthesis and biological function in mammals. Glycobiology. 2003;13(7):41R–53R. Epub 20030319. doi: 10.1093/glycob/cwg054. PubMed PMID: 12651883.

12. Majima Y, Harada T, Shimizu T, Takeuchi K, Sakakura Y, Yasuoka S, et al. Effect of biochemical components on rheologic properties of nasal mucus in chronic sinusitis. American journal of respiratory and critical care medicine. 1999;160(2):421–6. Epub 1999/08/03. PubMed PMID: 10430708.

13. Innes AL, McGrath KW, Dougherty RH, McCulloch CE, Woodruff PG, Seibold MA, et al. The H antigen at epithelial surfaces is associated with susceptibility to asthma exacerbation. American journal of respiratory and critical care medicine. 2011;183(2):189–94. Epub 2010/08/25. doi: 10.1164/rccm.201003-0488OC. PubMed PMID: 20732988; PubMed Central PMCID: 3040389.

14. Domino SE, Zhang L, Gillespie PJ, Saunders TL, Lowe JB. Deficiency of reproductive tract alpha(1,2)fucosylated glycans and normal fertility in mice with targeted deletions of the FUT1 or FUT2 alpha(1,2)fucosyltransferase locus. Mol Cell Biol. 2001;21(24):8336–45. Epub 2001/11/20. doi: 10.1128/MCB.21.24.8336-8345.2001. PubMed PMID: 11713270; PubMed Central PMCID: 99998.

15. Evans CM, Raclawska DS, Ttofali F, Liptzin DR, Fletcher AA, Harper DN, et al. The polymeric mucin Muc5ac is required for allergic airway hyperreactivity. Nature communications. 2015;6:6281. Epub 2015/02/18. doi: 10.1038/ncomms7281. PubMed PMID: 25687754; PubMed Central PMCID: 4333679.

16. Pilobello KT, Agrawal P, Rouse R, Mahal LK. Advances in lectin microarray technology: optimized protocols for piezoelectric print conditions. Curr Protoc Chem Biol. 2013;5(1):1–23. doi: 10.1002/9780470559277.ch120035. PubMed PMID: 23788322; PubMed Central PMCID: PMC4734107.

17. Piccotti L, Dickey BF, Evans CM. Assessment of intracellular mucin content in vivo. Methods in molecular biology. 2012;842:279–95. Epub 2012/01/20. doi: 10.1007/978-1-61779-513-8_17. PubMed PMID: 22259143.

18. Morgan LE, Jaramillo AM, Shenoy SK, Raclawska D, Emezienna NA, Richardson VL, et al. Disulfide disruption reverses mucus dysfunction in allergic airway disease. Nature communications. 2021;12(1):249. Epub 2021/01/13. doi: 10.1038/s41467-020-20499-0. PubMed PMID: 33431872; PubMed Central PMCID: PMC7801631.

19. Zhu Y, Ehre C, Abdullah LH, Sheehan JK, Roy M, Evans CM, et al. Munc13-2-/-baseline secretion defect reveals source of oligomeric mucins in mouse airways. J Physiol. 2008;586(7):1977–92. PubMed PMID: 18258655.

20. Evans CM, Williams OW, Tuvim MJ, Nigam R, Mixides GP, Blackburn MR, et al. Mucin is produced by clara cells in the proximal airways of antigen-challenged mice. Am JRespirCell MolBiol. 2004;31(4):382–94. PubMed PMID: 15191915.

21. Saku A, Hirose K, Ito T, Iwata A, Sato T, Kaji H, et al. Fucosyltransferase 2 induces lung epithelial fucosylation and exacerbates house dust mite-induced airway inflammation. The Journal of allergy and clinical immunology. 2019;144(3):698–709 e9. Epub 20190521. doi: 10.1016/j.jaci.2019.05.010. PubMed PMID: 31125592.

22. Alvarez CA, Qian E, Glendenning LM, Reynero KM, Kukan EN, Cobb BA. Acute and chronic lung inflammation drives changes in epithelial glycans. Front Immunol. 2023;14:1167908. Epub 20230522. doi: 10.3389/fimmu.2023.1167908. PubMed PMID: 37283757; PubMed Central PMCID: PMC10239862.

23. Warren KJ, Dickinson JD, Nelson AJ, Wyatt TA, Romberger DJ, Poole JA. Ovalbumin-sensitized mice have altered airway inflammation to agriculture organic dust. Respiratory research. 2019;20(1):51. Epub 20190307. doi: 10.1186/s12931-019-1015-0. PubMed PMID: 30845921; PubMed Central PMCID: PMC6407255.

24. Packer RJ, Shrine N, Hall R, Melbourne CA, Thompson R, Williams AT, et al. Genome-wide association study of chronic sputum production implicates loci involved in mucus production and infection. The European respiratory journal. 2023;61(6). Epub 20230615. doi: 10.1183/13993003.01667-2022. PubMed PMID: 37263751; PubMed Central PMCID: PMC10284065.

25. Ali S, Niang MA, N’Doye I, Critchlow CW, Hawes SE, Hill AV, et al. Secretor polymorphism and human immunodeficiency virus infection in Senegalese women. J Infect Dis. 2000;181(2):737–9. doi: 10.1086/315234. PubMed PMID: 10669366.

26. Kindberg E, Hejdeman B, Bratt G, Wahren B, Lindblom B, Hinkula J, et al. A nonsense mutation (428G-->A) in the fucosyltransferase FUT2 gene affects the progression of HIV-1 infection. AIDS. 2006;20(5):685–9. doi: 10.1097/01.aids.0000216368.23325.bc. PubMed PMID: 16514298.

27. Thorven M, Grahn A, Hedlund KO, Johansson H, Wahlfrid C, Larson G, et al. A homozygous nonsense mutation (428G-->A) in the human secretor (FUT2) gene provides resistance to symptomatic norovirus (GGII) infections. Journal of virology. 2005;79(24):15351–5. doi: 10.1128/JVI.79.24.15351-15355.2005. PubMed PMID: 16306606; PubMed Central PMCID: PMC1315998.

28. Imbert-Marcille BM, Barbe L, Dupe M, Le Moullac-Vaidye B, Besse B, Peltier C, et al. A FUT2 gene common polymorphism determines resistance to rotavirus A of the P[8] genotype. J Infect Dis. 2014;209(8):1227–30. Epub 20131125. doi: 10.1093/infdis/jit655. PubMed PMID: 24277741.

29. Santos-Cortez RLP, Chiong CM, Frank DN, Ryan AF, Giese APJ, Bootpetch Roberts T, et al. FUT2 Variants Confer Susceptibility to Familial Otitis Media. American journal of human genetics. 2018;103(5):679–90. Epub 20181025. doi: 10.1016/j.ajhg.2018.09.010. PubMed PMID: 30401457; PubMed Central PMCID: PMC6217759.

30. Taylor SL, Woodman RJ, Chen AC, Burr LD, Gordon DL, McGuckin MA, et al. FUT2 genotype influences lung function, exacerbation frequency and airway microbiota in non-CF bronchiectasis. Thorax. 2017;72(4):304–10. Epub 20160808. doi: 10.1136/thoraxjnl-2016-208775. PubMed PMID: 27503233.

31. Kashyap PC, Marcobal A, Ursell LK, Smits SA, Sonnenburg ED, Costello EK, et al. Genetically dictated change in host mucus carbohydrate landscape exerts a diet-dependent effect on the gut microbiota. Proceedings of the National Academy of Sciences of the United States of America. 2013;110(42):17059–64. Epub 20130923. doi: 10.1073/pnas.1306070110. PubMed PMID: 24062455; PubMed Central PMCID: PMC3800993.

32. Goto Y, Obata T, Kunisawa J, Sato S, Ivanov, II, Lamichhane A, et al. Innate lymphoid cells regulate intestinal epithelial cell glycosylation. Science. 2014;345(6202):1254009. Epub 20140821. doi: 10.1126/science.1254009. PubMed PMID: 25214634; PubMed Central PMCID: PMC4774895.

33. Domino SE, Hurd EA, Thomsson KA, Karnak DM, Holmen Larsson JM, Thomsson E, et al. Cervical mucins carry alpha(1,2)fucosylated glycans that partly protect from experimental vaginal candidiasis. Glycoconj J. 2009;26(9):1125–34. doi: 10.1007/s10719-009-9234-0. PubMed PMID: 19326211; PubMed Central PMCID: PMC2794911.

34. Kerr SC, Fischer GJ, Sinha M, McCabe O, Palmer JM, Choera T, et al. FleA Expression in Aspergillus fumigatus Is Recognized by Fucosylated Structures on Mucins and Macrophages to Prevent Lung Infection. PLoS Pathog. 2016;12(4):e1005555. Epub 2016/04/09. doi: 10.1371/journal.ppat.1005555. PubMed PMID: 27058347; PubMed Central PMCID: 4825926.

35. Houser J, Komarek J, Kostlanova N, Cioci G, Varrot A, Kerr SC, et al. A soluble fucose-specific lectin from Aspergillus fumigatus conidia--structure, specificity and possible role in fungal pathogenicity. PloS one. 2013;8(12):e83077. Epub 20131210. doi: 10.1371/journal.pone.0083077. PubMed PMID: 24340081; PubMed Central PMCID: PMC3858362.

36. Ishida H, Moritani T, Hata Y, Kawato A, Suginami K, Abe Y, et al. Molecular cloning and overexpression of fleA gene encoding a fucose-specific lectin of Aspergillus oryzae. Biosci Biotechnol Biochem. 2002;66(5):1002–8. doi: 10.1271/bbb.66.1002. PubMed PMID: 12092808.

37. Welsh KG, Rousseau K, Fisher G, Bonser LR, Bradding P, Brightling CE, et al. MUC5AC and a Glycosylated Variant of MUC5B Alter Mucin Composition in Children With Acute Asthma. Chest. 2017;152(4):771–9. Epub 2017/07/19. doi: 10.1016/j.chest.2017.07.001. PubMed PMID: 28716644; PubMed Central PMCID: 5624091.

38. Kirkham S, Sheehan JK, Knight D, Richardson PS, Thornton DJ. Heterogeneity of airways mucus: variations in the amounts and glycoforms of the major oligomeric mucins MUC5AC and MUC5B. BiochemJ. 2002;361(Pt 3):537–46. PubMed PMID: 11802783.

39. Sheehan JK, Howard M, Richardson PS, Longwill T, Thornton DJ. Physical characterization of a low-charge glycoform of the MUC5B mucin comprising the gel-phase of an asthmatic respiratory mucous plug. The Biochemical journal. 1999;338 (Pt 2):507–13. Epub 1999/02/20. PubMed PMID: 10024529; PubMed Central PMCID: 1220079.

40. Sheehan JK, Richardson PS, Fung DC, Howard M, Thornton DJ. Analysis of respiratory mucus glycoproteins in asthma: a detailed study from a patient who died in status asthmaticus. AmJ RespirCell MolBiol. 1995;13(6):748–56. PubMed PMID: 7576713.

41. Boruah BM, Kadirvelraj R, Liu L, Ramiah A, Li C, Zong G, et al. Characterizing human alpha-1,6-fucosyltransferase (FUT8) substrate specificity and structural similarities with related fucosyltransferases. The Journal of biological chemistry. 2020;295(50):17027–45. Epub 20201001. doi: 10.1074/jbc.RA120.014625. PubMed PMID: 33004438; PubMed Central PMCID: PMC7863877.

42. Jarva MA, Dramicanin M, Lingford JP, Mao R, John A, Jarman KE, et al. Structural basis of substrate recognition and catalysis by fucosyltransferase 8. The Journal of biological chemistry. 2020;295(19):6677–88. Epub 20200327. doi: 10.1074/jbc.RA120.013291. PubMed PMID: 32220931; PubMed Central PMCID: PMC7212660.

43. Garcia-Garcia A, Ceballos-Laita L, Serna S, Artschwager R, Reichardt NC, Corzana F, et al. Structural basis for substrate specificity and catalysis of alpha1,6-fucosyltransferase. Nature communications. 2020;11(1):973. Epub 20200220. doi: 10.1038/s41467-020-14794-z. PubMed PMID: 32080177; PubMed Central PMCID: PMC7033129.

44. Ihara H, Ikeda Y, Toma S, Wang X, Suzuki T, Gu J, et al. Crystal structure of mammalian alpha1,6-fucosyltransferase, FUT8. Glycobiology. 2007;17(5):455–66. Epub 20061215. doi: 10.1093/glycob/cwl079. PubMed PMID: 17172260.

45. Kadirvelraj R, Boruah BM, Wang S, Chapla D, Huang C, Ramiah A, et al. Structural basis for Lewis antigen synthesis by the alpha1,3-fucosyltransferase FUT9. Nat Chem Biol. 2023;19(8):1022–30. Epub 20230518. doi: 10.1038/s41589-023-01345-y. PubMed PMID: 37202521; PubMed Central PMCID: PMC10726971.

46. Pijnenborg JFA, Rossing E, Merx J, Noga MJ, Titulaer WHC, Eerden N, et al. Fluorinated rhamnosides inhibit cellular fucosylation. Nature communications. 2021;12(1):7024. Epub 20211202. doi: 10.1038/s41467-021-27355-9. PubMed PMID: 34857733; PubMed Central PMCID: PMC8640046.

47. Zafar H, Atif M, Atia Tul W, Choudhary MI. Fucosyltransferase 2 inhibitors: Identification via docking and STD-NMR studies. PloS one. 2021;16(10):e0257623. Epub 20211014. doi: 10.1371/journal.pone.0257623. PubMed PMID: 34648519; PubMed Central PMCID: PMC8516197.

