## supplement for "Requirement for Fucosyltransferase 2 in Allergic Airway Hyperreactivity and Mucus Obstruction"

### ONLINE SUPPLEMENT

Naoko Hara[1], Dorota S. Raclawska[1], Leslie E. Morgan[1], James C. NeeDell[1], Lucie Dao[1], Ayako Kato[1], Ana M. Jaramillo[1], Patrick S. Hume[2], Fernando Holguin[1], William J. Janssen[2], Eszter K. Vladar[1], Christopher M. Evans[1,3]*

[1]Division of Pulmonary Sciences and Critical Care Medicine, Department of Medicine, University of Colorado School of Medicine, Aurora, Colorado, USA

[2]Division of Pulmonary, Critical Care, and Sleep Medicine, Department of Medicine, National Jewish Health, Denver, Colorado, USA

[3]Research Service, Rocky Mountain Regional Veterans Affairs Medical Center, Aurora, Colorado, USA

#### Supplemental Methods

##### Human asthma tissue

Lung samples were obtained from deidentified tissue donated to National Jewish Health with Institutional Review Board approval. Briefly, tissue was screened and accepted for research from a 58 year-old female organ donor who was hospitalized for severe asthma leading to death. Other than the required diagnosis of asthma and a history of asthma since childhood, there were no additional diagnoses and no evidence of bacterial pneumonia. Following organ recovery, tissue was transported on ice and distributed for research studies in under 24 hours from the time of procurement. Tissues were fixed in 10% formalin, dissected into ~1 $cm^3$ pieces, embedded in paraffin, and sectioned into 5 µm thick histologic specimens.

##### Experimental animals

Studies were performed with approval of the animal care and use committee of the University of Colorado. Mice were housed under specific pathogen-free conditions in ventilated cages (≤5 mice per cage). Housing rooms were maintained at 22°C, 30-40% humidity, and a 14/10 (h/h) light/dark cycle with at least 12 fresh-air changes per h.

*Fut2*$^{-/-}$ mice were generated previously and were provided by Dr. Justin Sonnenburg at Stanford University (1). This line was produced on a mixed 129X1/SvJ and C57BL/6J background and was crossed to a C57BL/6J congenic background for >12 generations. The line was maintained at the University of Colorado by routinely backcrossing to C57BL/6J mice purchased from the Jackson Laboratories (Bar Harbor, ME). Experimental animals were produced by intercrossing heterozygous *Fut2*$^{+/-}$ animals and by crossing *Fut2*$^{+/+}$ or *Fut2*$^{-/-}$ homozygous pairs. Mice were placed into experimental and control groups chronologically. Males and females were used starting at 7-8 wks age. Sex distributions are displayed in results using square symbols for males and circles for females.

**Allergic lung inflammation modeling**

Mice were challenged using aerosolized *Aspergillus oryzae* extract (2) (AOE; Cat. P6110, Sigma, St. Louis, MO) administered weekly for a total of 4 exposures. Endpoint analyses were studied 48 h after the last AOE challenge. For each exposure, a 5 ml volume of a 10% v/v solution of AOE in PBS was delivered via an Ultravent jet nebulizer (Covidien, Dublin, Ireland) operating at 30 psi to produce respirable particles via nose-only inhalation (3).

**Mucin detection**

To detect human mucins by immunofluorescence, triple labeling was performed to detect MUC5AC, MUC5B, and Fuc(α1-2)Gal in the airways. Dewaxed tissues underwent antigen retrieval in 10 mM citrate buffer. After incubation with Carbo-Free Blocking Solution (Vector Laboratories, Cat. SP-5040-125), a mixture containing mouse-anti-human MUC5AC (clone 45M1, Novus, Cat. NBP2-15196, 1:1000 dilution), rabbit-anti-human MUC5B (Sigma, Cat. HPA008246, 1:500 dilution), and biotin-conjugated *Ulex europaeus* agglutinin-1 (UEA1, Vector, B-1065-2, 8 μg/ml) was applied. UEA1 is a lectin that is selective for detection of the glycan structure Fuc(α1-2)Gal (4), which is abbreviated as α-Fuc hereafter. After washing in PBS containing 0.1% Tween 20, Alexa 546-conjugated goat-anti-mouse IgG, Alexa 488-conjugated goat-anti-rabbit IgG, and Alexa 647-conjugated streptavidin were diluted together applied for detection. Slides were washed and then cover-slipped with fluorescent mounting media containing DAPI. Microscopic imaging was performed using an Olympus BX63 microscope and cellSens software (Olympus USA, Center Valley, PA).

Mouse mucins were detected using affinity purified polyclonal rabbit-anti-mouse MUC5AC produced against the peptide N-CHALGDTSHAESSEQEFKSKESEEHGQQLAFR (Pacific Immunology, Ramona, CA, 1:2500 dilution), monoclonal mouse-anti-mouse MUC5B (clone

MDA-3E1, Cat. MABT899, MilliporeSigma, Burlington, MA, 1:500 dilution), and UEA1 (as above). For microscopy, Alexa 546-conjugated goat-anti-rabbit IgG, Alexa 488-conjugated goat-anti-mouse IgG, and Alexa 647-conjugated streptavidin, and DAPI were applied.

**Assessment of lung inflammation**

Allergic inflammation was confirmed by enumeration of leukocytes in lung lavage and mediators in lung tissues. Lung lavage was performed by instilling 5 x 0.5 ml PBS into the right lungs of euthanized, tracheostomized mice. To improve homogeneous sampling on the first round of lavage, the same 0.5 ml of PBS was instilled and retrieved 3 times and then collected in a 1.5 ml tube. Aliquots of unmanipulated "neat" first-round lavage fluid were stored at -20° C for subsequent immunoblot analyses. Due to the sizes of mucin glycopolymers, neat samples are required as material will sediment into pellets upon centrifugation using standard lung lavage processing procedures. Remaining portions of first lavage and subsequent 2.0 ml collections were counted on a hemacytometer and on Cytospin slides stained with Giemsa dye.

After lavage, right lung lobes were excised, homogenized, aliquotted, and frozen on dry ice. Portions of homogenates (~50 mg per mouse) were used for multiplex analysis of the cytokines IFN-γ, IL-1β, IL-2, IL-4, IL-5, IL-6, IL-9, IL-10, IL-12p70 (IL-12), IL-15, IL-17A/F, IL-27p28 (IL-30), IL-33, and TNF and the chemokines IP-10 (CXCL10), KC/GRO (CXCL1), MCP-1 (CCL2), MIP-2 (CXCL2) , and MIP-1α (CCL3) using a V-PLEX Mouse Cytokine 19-Plex Kit (Mesoscale Discovery, Rockville, MD).

Briefly, lung homogenates were thawed and denatured in RIPA buffer (0.1% SDS w/v, 1% Triton-X100 v/v, 1 mM DTT, 3 M Urea, 10x protease inhibitor cocktail, Sigma). After centrifugation to remove debris (14,000 x g, 10 min, 4° C), total protein concentrations were assessed using a bicinchoninic acid (BCA) assay (ThermoFisher). For V-PLEX assays, 50 μl samples were applied at ~2-8 μg/μl total protein concentrations per the manufacturer's instructions. Cytokine concentrations were normalized to total protein levels determined by BCA assay. Individual datapoints are shown in Table E1.

**Mucin glycoprotein blotting**

Neat lavage samples were thawed, and 27 μl aliquots were mixed with 3 μl of non-reducing loading buffer and then loaded into 1% SDS agarose gels. After electrophoresis (100 V, 2 h), samples were transfer to PVDF membranes by vacuum blot (5, 6). Membranes were blocked with Carbo-Free Blocking Solution (Vector), and mucins were probed with mixtures of UEA1 (1:1,000, 2 μg/ml) and either polyclonal rabbit-anti-mouse Muc5ac (1:2,000) or rabbit-anti-mouse Muc5b (1:5,000) (7, 8). Labels were detected with IRDye 680RD-conjugated streptavidin and IRDye 800CW-conjugated goat anti-rabbit IgG (LI-COR, Lincoln, NE, diluted 1:15,000). Imaging was performed on an Odyssey CLx, and analysis was performed using Image Studio software (LI-COR). Since both anti-mucin antisera labels were generated in rabbits, blotting was performed using identical samples loaded in wells on separate halves of the same agarose gels. After vacuum transfer, PVDF membranes were cut, and the steps above were performed in parallel.

**Airway hyperreactivity**

As reported previously (2), lung function was measured 48 h after the last AOE challenge using a flexiVent (Scireq, Montreal, Quebec, Canada). Mice were anesthetized with urethane (2.0 g/kg, i.p.), tracheostomized with a blunt beveled 18 G Luer stub, and ventilated (150 breaths/min, 10 ml/kg, against 3 cm $H_2O$ positive end expiratory pressure with paralysis maintained continuous infusion of succinylcholine chloride (10 μg/g/min, i.p.).

Total lung resistance ($R_L$) and airway resistance ($R_{AW}$) were determined at baseline and in response to successive, increasing doses of methacholine chloride (MCh; Sigma, St. Louis, MO, Cat no. A2251; 0.1-10 mg/ml) administered via an in-line ultrasonic nebulizer. During peak bronchoconstriction in response to 10 mg/ml MCh, mice were given anesthetic overdose (6 g/kg, i.p.), ventilation was stopped, and lungs were fixed *in situ*.

**Mucus plugging**

Tissues were fixed using a method described previously (2, 6). Briefly, while kept and at end tidal volume, the lower sternum was exposed, and 400 µl of methacarn was injected into the pleural space using a 36 G needle. After fixation *in situ* the trachea was ligated, and the lungs were removed and fixed in methacarn at 4° C overnight. Fixed lungs were transferred to absolute methanol, and total lung volume was calculated using displacement of absolute methanol. Lungs were then cut into 2-3 mm cubes (~30 per lung), paraffin embedded in random orientations, sectioned at 5 µm thickness, and collected on glass slides for histologic assessments.

Mucins were labeled using alcian blue-periodic acid Schiff's (AB-PAS) stain (9), and slides were scanned under brightfield illumination using a 2x objective. Each tissue sample was then imaged from its center using a 10x objective. Total airway vs. parenchyma fractions were counted using a 250 x 250 µm grid, and airway mucin volume fractions were quantified using a 50 x 50 µm grid. Volume fractions were then converted to volumes using displacement measurements.

**Statistical analyses**

All statistical analyses were performed using GraphPad Prism 10.2.2 (397). Inflammation, AHR, and mucin secretion data were compared between groups using unpaired two-way student's *t*-

tests or Mann-Whitney U-tests where appropriate. For multiple comparisons, ANOVA with a Dunnett post-hoc correction or a Kruskal-Wallis test with Dunn's post-hoc correction was used. For dose response tests, curves were generated, and non-linear fit regression was used to calculate best-fit slopes of semi-log data. Slopes were then compared using ANOVA as described above. Sample sizes were chosen based on the numbers of WT saline vs. allergic mice required to achieve statistically significant difference. $R_{AW}$ data for one AOE challenged *Fut*$^{+/+}$ mouse was removed from analyses due to poor fitting on input impedance models, but its $R_L$ companion point was valid and was thus included in final analyses.

#### Supplemental References

1. Domino SE, Zhang L, Gillespie PJ, Saunders TL, Lowe JB. Deficiency of reproductive tract alpha(1,2)fucosylated glycans and normal fertility in mice with targeted deletions of the FUT1 or FUT2 alpha(1,2)fucosyltransferase locus. Mol Cell Biol. 2001;21(24):8336-45. Epub 2001/11/20. doi: 10.1128/MCB.21.24.8336-8345.2001. PubMed PMID: 11713270; PubMed Central PMCID: 99998.

2. Evans CM, Raclawska DS, Ttofali F, Liptzin DR, Fletcher AA, Harper DN, et al. The polymeric mucin Muc5ac is required for allergic airway hyperreactivity. Nature communications. 2015;6:6281. Epub 2015/02/18. doi: 10.1038/ncomms7281. PubMed PMID: 25687754; PubMed Central PMCID: 4333679.

3. Foster MW, Yang Z, Potts EN, Michael Foster W, Que LG. S-nitrosoglutathione supplementation to ovalbumin-sensitized and -challenged mice ameliorates methacholine-induced bronchoconstriction. American journal of physiology Lung cellular and molecular physiology. 2011;301(5):L739-44. Epub 2011/07/26. doi: 10.1152/ajplung.00134.2011. PubMed PMID: 21784966; PubMed Central PMCID: 3213990.

4. Pilobello KT, Agrawal P, Rouse R, Mahal LK. Advances in lectin microarray technology: optimized protocols for piezoelectric print conditions. Curr Protoc Chem Biol. 2013;5(1):1-23. doi: 10.1002/9780470559277.ch120035. PubMed PMID: 23788322; PubMed Central PMCID: PMC4734107.

5. Piccotti L, Dickey BF, Evans CM. Assessment of intracellular mucin content in vivo. Methods in molecular biology. 2012;842:279-95. Epub 2012/01/20. doi: 10.1007/978-1-61779-513-8_17. PubMed PMID: 22259143.

6. Morgan LE, Jaramillo AM, Shenoy SK, Raclawska D, Emezienna NA, Richardson VL, et al. Disulfide disruption reverses mucus dysfunction in allergic airway disease. Nature communications. 2021;12(1):249. Epub 2021/01/13. doi: 10.1038/s41467-020-20499-0. PubMed PMID: 33431872; PubMed Central PMCID: PMC7801631.

7. Zhu Y, Ehre C, Abdullah LH, Sheehan JK, Roy M, Evans CM, et al. Munc13-2-/- baseline secretion defect reveals source of oligomeric mucins in mouse airways. J Physiol. 2008;586(7):1977-92. PubMed PMID: 18258655.

8. Roy MG, Livraghi-Butrico A, Fletcher AA, McElwee MM, Evans SE, Boerner RM, et al. Muc5b is required for airway defence. Nature. 2014;505(7483):412-6. Epub 2013/12/10. doi: 10.1038/nature12807. PubMed PMID: 24317696; PubMed Central PMCID: 4001806.

9. Evans CM, Williams OW, Tuvim MJ, Nigam R, Mixides GP, Blackburn MR, et al. Mucin is produced by clara cells in the proximal airways of antigen-challenged mice. Am JRespirCell MolBiol. 2004;31(4):382-94. PubMed PMID: 15191915.

**Table E1. Individual values of mouse cytokine panel results (µg analyte per µg total protein).**

| | | Fut2+/+ Sal | | | Fut2-/- Sal | | | Fut2+/+ AOE | | | | | | Fut2-/- AOE | | | |
|---|---|---|---|---|---|---|---|---|---|---|---|---|---|---|---|---|---|
| | **Mouse ID** | **210** | **247** | **496** | **204** | **491** | **493** | **161** | **206** | **239** | **243** | **345** | **489** | **246** | **483** | **485** | **492** |
| **Analyte** | IL-1β | 0.70 | 0.46 | 0.16 | 0.34 | 0.43 | 1.74 | 3.94 | 2.14 | 5.03 | 9.42 | 2.32 | 4.01 | 3.41 | 1.67 | 1.85 | 1.30 |
| | IL-6 | 5.00 | 4.45 | 2.62 | 3.79 | 3.89 | 6.10 | 43.27 | 12.14 | 10.57 | 29.12 | 13.53 | 15.19 | 79.77 | 28.11 | 20.84 | 20.84 |
| | TNF | 0.53 | 0.35 | 0.27 | 0.38 | 0.37 | 1.01 | 1.45 | 1.02 | 0.65 | 0.64 | 0.67 | 1.02 | 0.52 | 1.00 | 0.83 | 0.95 |
| | IFNγ | 1.22 | 0.24 | 0.25 | 0.43 | 0.44 | 0.46 | 0.47 | 0.46 | 0.18 | 0.14 | 0.29 | 0.19 | 0.13 | 0.26 | 0.24 | 0.56 |
| | IL-12p70 (IL-12) | 9.49 | 5.53 | 4.25 | 6.10 | 6.50 | 9.04 | 15.15 | 7.25 | 5.15 | 4.53 | 3.78 | 7.83 | 3.85 | 6.04 | 5.96 | 4.44 |
| | IL-4 | 0.14 | 0.07 | 0.06 | 0.11 | 0.12 | 0.16 | 1.29 | 0.51 | 0.45 | 1.25 | 1.86 | 2.06 | 1.27 | 2.79 | 2.73 | 3.50 |
| | IL-5 | 0.53 | 0.39 | 0.25 | 0.38 | 0.37 | 0.46 | 2.18 | 3.14 | 0.65 | 0.73 | 1.60 | 1.47 | 0.91 | 2.99 | 2.49 | 3.55 |
| | IL-9 | 0.76 | 0.51 | 0.34 | 0.33 | 0.39 | 0.86 | 1.83 | 2.10 | 0.90 | 1.06 | 0.65 | 0.96 | 0.68 | 0.91 | 0.71 | 0.88 |
| | IL-17 | 4.90 | 4.36 | 2.57 | 3.72 | 3.82 | 5.98 | 42.43 | 11.90 | 10.36 | 28.55 | 13.27 | 14.90 | 78.23 | 27.57 | 20.44 | 20.44 |
| | IL-2 | 0.34 | 0.23 | 0.15 | 0.15 | 0.18 | 0.39 | 0.82 | 0.95 | 0.41 | 0.48 | 0.29 | 0.43 | 0.31 | 0.41 | 0.32 | 0.40 |
| | IL-10 | 0.42 | 0.40 | 0.29 | 0.41 | 0.40 | 0.33 | 1.02 | 0.51 | 0.44 | 0.29 | 0.37 | 0.52 | 0.31 | 0.52 | 0.39 | 0.41 |
| | IL-15 | 46.9 | 34.5 | 22.6 | 33.4 | 32.7 | 41.0 | 194.0 | 279.1 | 58.1 | 64.6 | 142.4 | 130.9 | 81.2 | 265.9 | 220.8 | 315.5 |
| | IL-27p28 (IL-30) | 1.14 | 0.57 | 0.49 | 0.88 | 0.95 | 1.31 | 10.38 | 4.10 | 3.57 | 10.05 | 14.93 | 16.56 | 10.21 | 22.41 | 21.88 | 28.06 |
| | IL-33 | 10.34 | 2.01 | 2.12 | 3.62 | 3.76 | 3.86 | 3.99 | 3.89 | 1.49 | 1.19 | 2.47 | 1.60 | 1.13 | 2.23 | 2.01 | 4.74 |
| | CXCL1 (KC) | 3.19 | 0.74 | 0.83 | 1.35 | 1.55 | 12.16 | 5.63 | 1.54 | 1.47 | 2.63 | 1.21 | 1.87 | 3.38 | 1.46 | 1.09 | 4.67 |
| | CXCL2 (MIP-2) | 0.43 | 0.25 | 0.19 | 0.28 | 0.30 | 0.41 | 0.69 | 0.33 | 0.23 | 0.21 | 0.17 | 0.36 | 0.18 | 0.28 | 0.27 | 0.20 |
| | CXCL10 (IP-10) | 0.23 | 0.21 | 0.15 | 0.22 | 0.21 | 0.18 | 0.55 | 0.27 | 0.24 | 0.16 | 0.20 | 0.28 | 0.17 | 0.28 | 0.21 | 0.22 |
| | CCL2 (MCP-1) | 0.39 | 0.26 | 0.09 | 0.19 | 0.24 | 0.98 | 2.22 | 1.20 | 2.84 | 5.31 | 1.31 | 2.26 | 1.92 | 0.94 | 1.04 | 0.73 |
| | CCL3 (MIP-1α) | 1.60 | 0.37 | 0.42 | 0.67 | 0.77 | 6.08 | 2.81 | 0.77 | 0.73 | 1.31 | 0.60 | 0.93 | 1.69 | 0.73 | 0.54 | 2.33 |

**Table E2. Summary data of mouse cytokine panel results (µg analyte per µg total protein).**

| | Fut2+/+ Sal | | Fut2-/- Sal | | Fut2+/+ AOE | | Fut2-/- AOE | | Kruskal-Wallis | | | Mann-Whitney |
|---|---|---|---|---|---|---|---|---|---|---|---|---|
| Analyte | Mean | SEM | Mean | SEM | Mean | SEM | Mean | SEM | +/+ (Sal v. AOE) | -/- (Sal v. AOE) | AOE (+/+ v. -/-) | AOE (+/+ v. -/-) |
| IL-1β | 0.44 | 0.16 | 0.84 | 0.45 | 4.48 | 1.09 | 2.06 | 0.47 | **0.02*** | >0.99 | 0.77 | **0.04*** |
| IL-6 | 4.02 | 0.72 | 4.60 | 0.75 | 20.64 | 5.28 | 37.39 | 14.23 | 0.20 | 0.06 | >0.99 | 0.26 |
| TNF | 0.38 | 0.08 | 0.58 | 0.21 | 0.91 | 0.13 | 0.83 | 0.11 | 0.07 | >0.99 | >0.99 | 0.61 |
| IFNγ | 0.57 | 0.33 | 0.44 | 0.01 | 0.29 | 0.06 | 0.30 | 0.09 | >0.99 | >0.99 | >0.99 | >0.99 |
| IL-12p70 (IL-12) | 6.42 | 1.58 | 7.21 | 0.92 | 7.28 | 1.70 | 5.07 | 0.55 | >0.99 | 0.62 | >0.99 | 0.48 |
| IL-4 | 0.09 | 0.03 | 0.13 | 0.02 | 1.24 | 0.27 | 2.57 | 0.47 | 0.18 | 0.06 | >0.99 | >0.99 |
| IL-5 | 0.39 | 0.08 | 0.40 | 0.03 | 1.63 | 0.38 | 2.49 | 0.57 | 0.25 | **0.05*** | >0.99 | 0.26 |
| IL-9 | 0.54 | 0.12 | 0.53 | 0.17 | 1.25 | 0.24 | 0.80 | 0.06 | 0.14 | >0.99 | >0.99 | 0.17 |
| IL-17 | 3.95 | 0.70 | 4.51 | 0.74 | 20.24 | 5.18 | 36.67 | 13.95 | 0.20 | 0.06 | >0.99 | 0.26 |
| IL-2 | 0.24 | 0.05 | 0.24 | 0.08 | 0.56 | 0.11 | 0.36 | 0.03 | 0.14 | >0.99 | >0.99 | 0.17 |
| IL-10 | 0.37 | 0.04 | 0.38 | 0.02 | 0.53 | 0.11 | 0.41 | 0.04 | >0.99 | >0.99 | >0.99 | 0.76 |
| IL-15 | 34.65 | 7.02 | 35.73 | 2.66 | 144.8 | 33.96 | 220.8 | 50.41 | 0.25 | **0.05*** | >0.99 | 0.26 |
| IL-27p28 (IL-30) | 0.73 | 0.21 | 1.04 | 0.13 | 9.93 | 2.19 | 20.64 | 3.75 | 0.18 | 0.06 | >0.99 | 0.07 |
| IL-33 | 4.82 | 2.76 | 3.75 | 0.07 | 2.44 | 0.51 | 2.53 | 0.78 | >0.99 | >0.99 | >0.99 | >0.99 |
| CXCL1 (KC) | 1.59 | 0.80 | 5.02 | 3.57 | 2.39 | 0.68 | 2.65 | 0.84 | >0.99 | >0.99 | >0.99 | 0.91 |
| CXCL2 (MIP-2) | 0.29 | 0.07 | 0.33 | 0.04 | 0.33 | 0.08 | 0.23 | 0.03 | >0.99 | 0.62 | >0.99 | 0.48 |
| CXCL10 (IP-10) | 0.20 | 0.02 | 0.20 | 0.01 | 0.28 | 0.06 | 0.22 | 0.02 | >0.99 | >0.99 | >0.99 | 0.76 |
| CCL2 (MCP-1) | 0.25 | 0.09 | 0.47 | 0.25 | 2.52 | 0.61 | 1.16 | 0.26 | **0.02*** | >0.99 | 0.77 | **0.04*** |
| CCL3 (MIP-1α) | 0.79 | 0.40 | 2.51 | 1.78 | 1.19 | 0.34 | 1.32 | 0.42 | >0.99 | >0.99 | >0.99 | 0.91 |

*Significant difference ($p<0.05$).

**Supplemental Figure Legends**

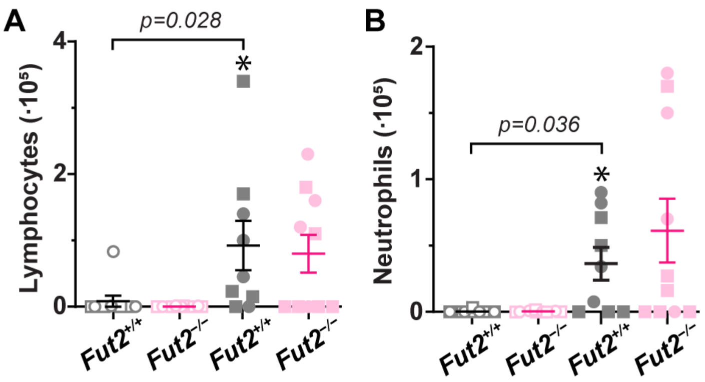

**Figure E1. Fut2 deficiency does not reduce inflammation after AOE challenge.** Lung lavage fluid was obtained from saline and AOE challenged *Fut2*$^{+/+}$ (gray-black) and *Fut2*$^{-/-}$ (magenta) mice. AOE challenge resulted in increases in lymphocytes and neutrophils in *Fut2*$^{+/+}$ mice, but not in *Fut2*$^{-/-}$ mice. Kruskal-Wallis ANOVA was used to compare AOE challenged *Fut2*$^{+/+}$ mice (filled gray shapes, n = 9 biological replicates), AOE challenged *Fut2*$^{-/-}$ mice (filled magenta shapes, n = 10 biological replicates), saline challenged *Fut2*$^{+/+}$ mice (open gray shapes, n = 10 biological replicates), and saline challenged *Fut2*$^{-/-}$ mice (open magenta shapes, n = 9 biological replicates). '*', $p < 0.05$ using Dunn's post-hoc test for multiple comparisons (p-values are shown). In AOE vs saline comparisons for *Fut2*$^{-/-}$ mice, p=0.47 for lymphocytes, and p=0.21 for neutrophils. Squares identify males, and circles identify females. No significant differences were observed using sex as a biological variable.

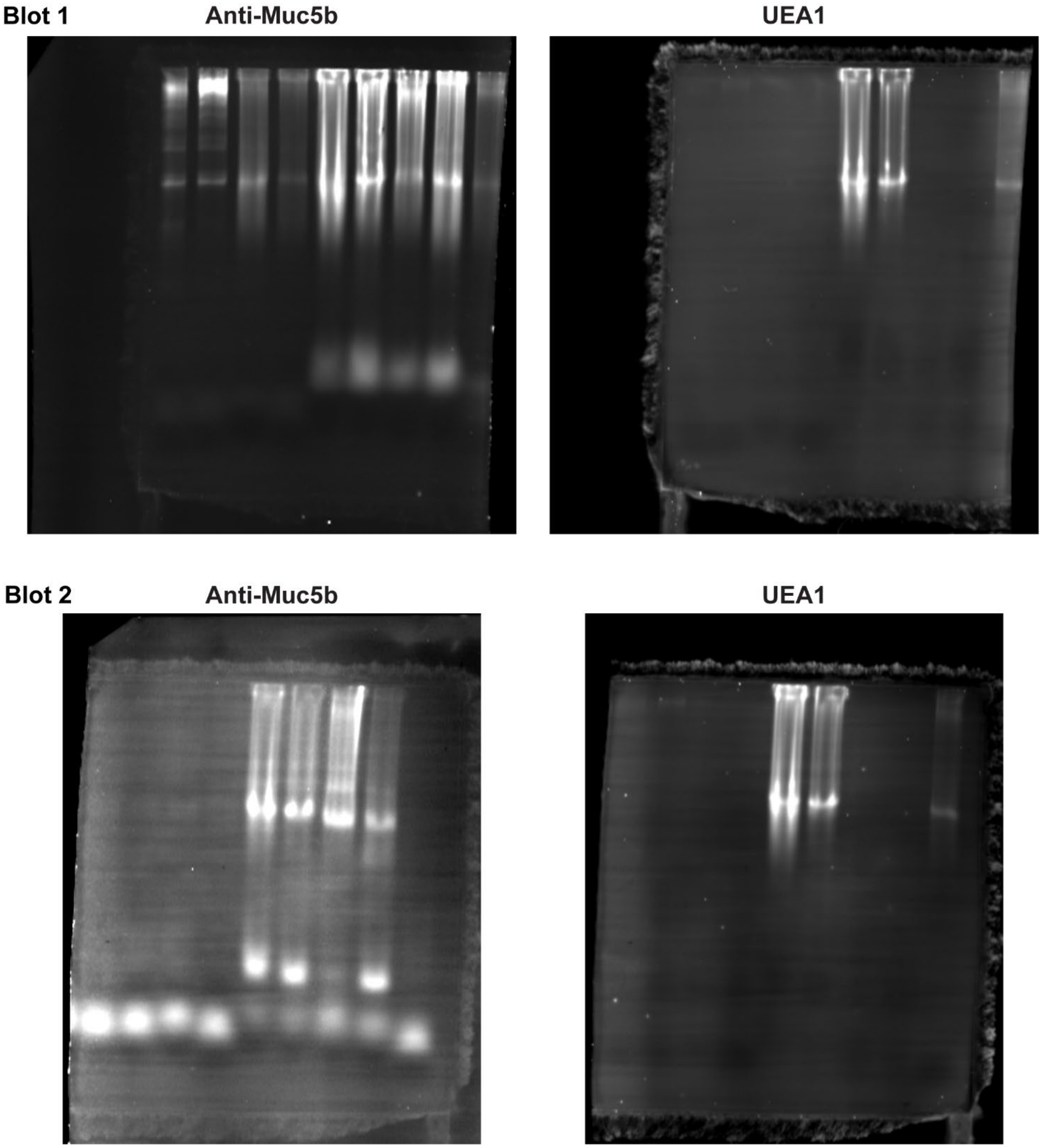

**Figure E2. Fut2 deficiency abolishes airway mucin fucosylation.** Combined immuno- and lectin blotting was performed on neat lavage. Equal volumes of lavage fluid (27 µl) were loaded per well, separated in 1% SDS agarose gels, and transferred to PVDF membranes. Membranes were probed with biotinylated UEA1 (1:1,000, 2 µg/ml) and either rabbit-anti-MUC5AC (1:2,000) or rabbit-anti-MUC5B (1:5,000). For secondary detection, Alexa 680-conjugated streptavidin and Alexa 800-conjugated labeled goat-anti-rabbit probes (Licor, 1:15,000) were applied. Monochrome images were acquired and pseudocolored magenta (MUC5AC), cyan (MUC5B), or yellow (UEA1). Images show original, uncropped monochrome versions of each layer from blots with anti-mucin antibodies and UEA1.

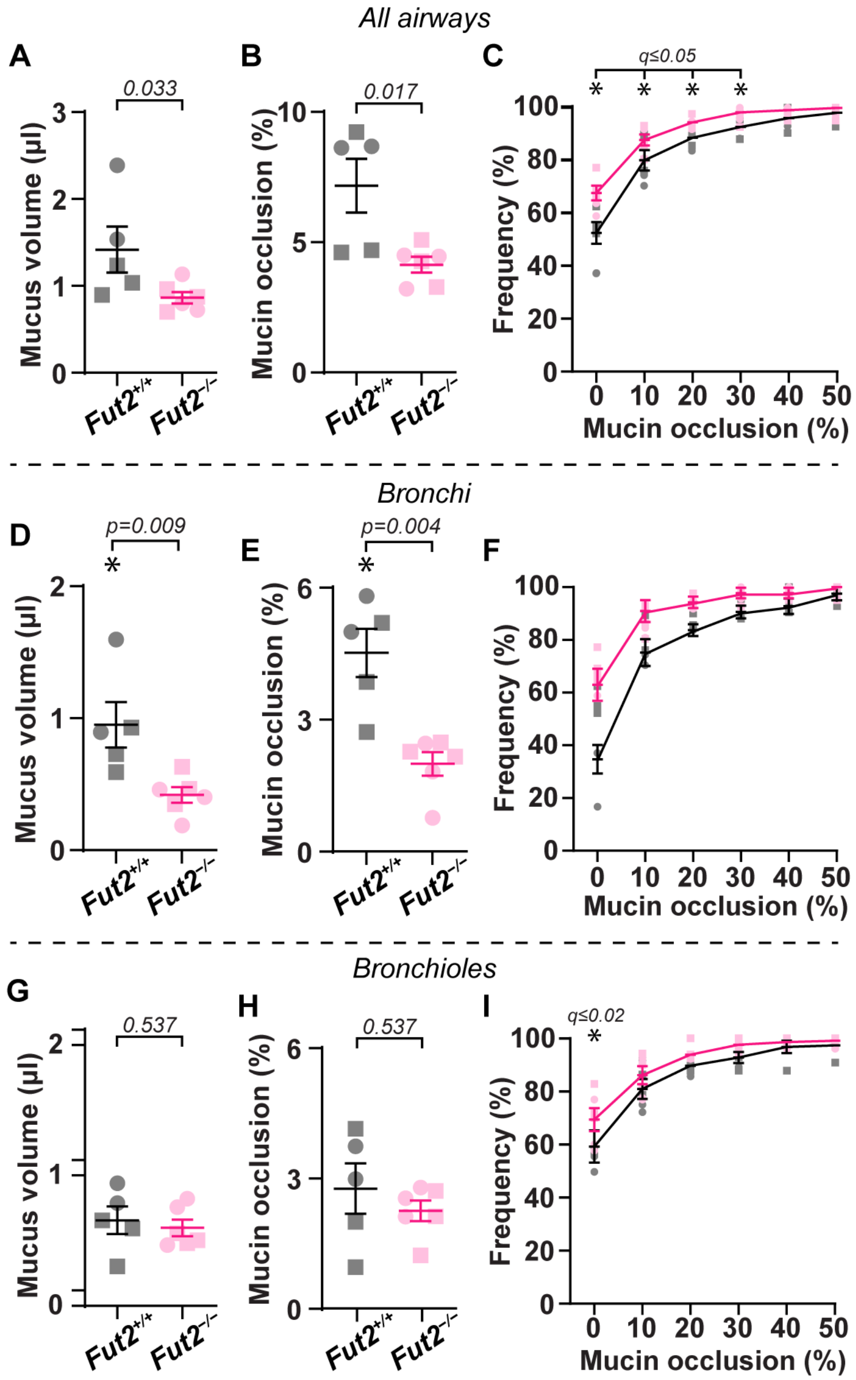

**Figure E3. Fut2 deficiency reduces airway mucus plugging.** After lung mechanics studies, lungs were fixed with methacarn to preserve mucus in airspaces. Data show summarized results from all airways (**A-C**), bronchial airways (**D-F**), and bronchioles (**G-I**). Calculated mucus volumes (**A,D,G**), fractional mucin occlusion (**B,E,H**), and heterogeneous plugging (**C,F,I**) are shown. Note: Data in D-F are identical to Figure 5 in the main manuscript. Data in **A, B, D, E, G**, and **H** are means ± sem for *Fut2*$^{+/+}$ (black lines, n = 5 mice) and *Fut2*$^{-/-}$ (magenta lines, n = 6 mice) on scatter plots. P-values are shown with '*' depicting p<0.05 by two-tailed Mann-Whitney tests. Data in **C**, **F**, and **I** are means ± sem displayed on cumulative frequency distribution graphs. Fractional occlusion means per animal were evaluated between *Fut2*$^{+/+}$ and *Fut2*$^{-/-}$ mice using multiple unpaired t-tests with two-stage step-up methods for multiple comparisons at a false discovery rate cut-off (q-value) set at 0.05. '*' demonstrates significance, and q-values are shown in **C**, **F**, and **I**. Semi-transparent shapes identify results from individual mice in **A, B, D, E, G**, and **H**. Squares identify males, and circles identify females. No significant differences were observed using sex as a biological variable. Data in **D-F** are identical to **Figure 7C-E**.
